# Integrating Molecular Simulation and Experimental Data: A Bayesian/Maximum Entropy reweighting approach

**DOI:** 10.1101/457952

**Authors:** Sandro Bottaro, Tone Bengtsen, Kresten Lindorff-Larsen

## Abstract

We describe a Bayesian/Maximum entropy (BME) procedure and software to construct a conformational ensemble of a biomolecular system by integrating molecular simulations and experimental data. First, an initial conformational ensemble is constructed using for example Molecular Dynamics or Monte Carlo simulations. Due to potential inaccuracies in the model and finite sampling effects, properties predicted from simulations may not agree with experimental data. In BME we use the experimental data to refine the simulation so that the new conformational ensemble has the following properties: (i) the calculated averages are close to the experimental values taking uncertainty into account and (ii) it maximizes the relative Shannon entropy with respect to the original simulation ensemble. The output of this procedure is a set of optimized weights that can be used to calculate arbitrary properties and distributions. Here, we provide a practical guide on how to obtain and use such weights, how to choose adjustable parameters and discuss shortcomings of the method.

## 1 Introduction

Experimental determination of biomolecular structure and dynamics is an important and difficult problem in molecular biology. A large variety of techniques to tackle this problem exists, including X-ray/neutron diffraction and scattering experiments, nuclear magnetic resonance (NMR) spectroscopy, cryo-electron microscopy, and a plethora of other techniques. These experiments often result in noisy and incomplete data, making it non-trivial to solve the inverse problem of reconstructing structural and dynamical molecular properties from experiments alone [1].

Computer simulations based e.g. on physics-derived or knowledge-based models can in principle provide a detailed thermodynamic description for arbitrary molecular systems. Performing such simulations is, however, often not sufficient, due to inaccuracies of the molecular models (force fields), and the high computational cost associated with extensive simulations.

For these reasons, there exists a number of integrative approaches in which simulations and experiments are combined [2–6]. In some of these approaches, a physical model (i.e. the force field) is complemented by a set of experimental restraints favouring molecular conformations that individually match experimental data. When studying flexible molecular systems that populate multiple conformations, however, this approach leads to wrong results, because it drives the simulation towards intermediates that are not representative of any of the relevant states (Fig. 1) [6–11]. Such problems may be particularly relevant when the systems are structurally very heterogeneous, and when the experimental measurements have nonlinear dependencies of the conformational properties.

**Fig. 1.**
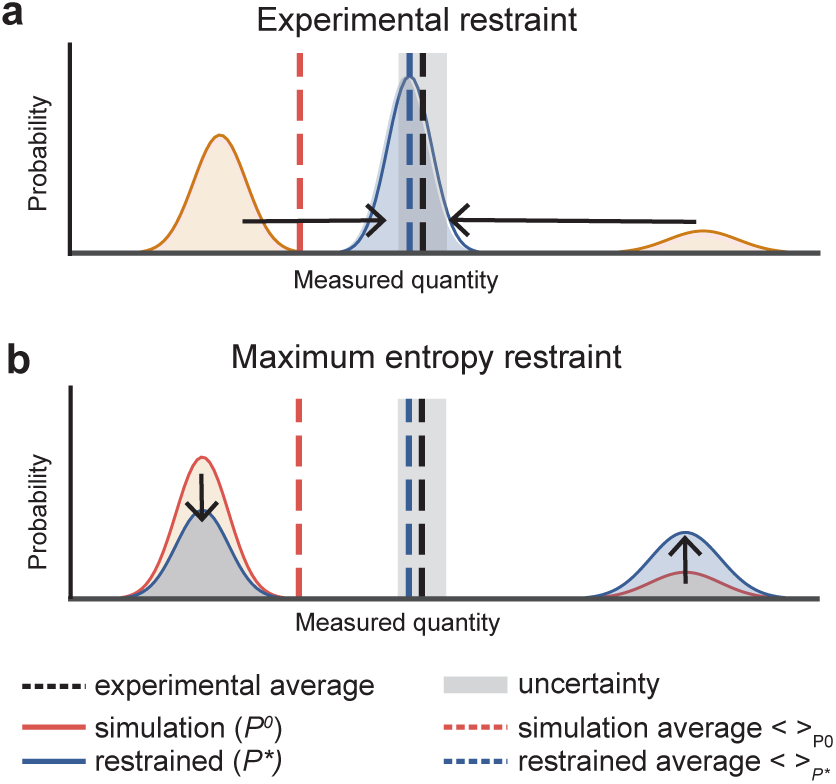
Schematic example showing two different strategies to restrain simulations using experimental data. When performing a molecular simulation, samples from a prior distribution, *P*^0^, are generated, using for example the Boltzmann distribution from a Molecular Dynamics (MD) simulation. When a calculated average 8·9*P*^0^ does not match the experimental measurement, it is possible to use the experimental data to modify the prior distribution, resulting in new, optimized probability distribution *P*\* (also called the posterior). The new average 8·9*P*\* matches the experimental data with some level of uncertainty. Different strategies to derive the posterior distribution are possible. a A common choice is to require all individual molecular conformations to match the experimental data within uncertainty. b In the ME formalism, *P*\* is instead the minimal modification to *P*^0^ that brings the calculated averages to match the experimental data, resulting in the optimal combination between simulations and experiments.

Maximum-entropy (ME) [12] approaches treat experimental data as time/ensemble averages, and make it possible to combine the physico-chemical information deriving from the simulation with experimental knowledge. In its basic implementation, however, the ME formalism does not take uncertainty and noise into account. Recently, it has, however, been shown how to generalize ME to take into account the uncertainty associated with the experimental data [10, 13, 14].

There are two principally different ways of combining the experimental data and molecular force field to generate the ME ensemble [11]. One set of methods uses the experimental data directly as restraints during the simulations, thus generating samples directly from the target probability distribution. An advantage of this approach is that one can focus sampling efforts only on the most relevant regions of conformational landscape, but comes at the cost of both additional complexity in simulation software, and a requirement that the experimental data can be calculated rapidly from molecular conformations and with analytical gradients. The second approach uses standard simulation methods to generate a conformational ensemble, which is then reweighted afterwards using the experimental data to generate a weighted ensemble representing the target probability distribution *P*\*. The advantages of this approach includes its simplicity, the fact that it can easily be combined with numerous methods for enhanced sampling, and that one can use rather complex models for calculating experimental observables [15]. It is this second method that is the focus on this paper.

Thus, we here describe a procedure to apply the ME approach to existing simulations sampled from some prior probability distribution. The procedure is in essence identical to the Bayesian inference of ensembles (BioEn) [10, 16], also originally called ensemble refinement of SAXS (EROS) [17, 18], that consists in finding the set of weights that maximize a functional that ranks configuration space distributions. Here, we explicitly make use of the ME formalism: this makes it possible to simplify considerably the minimization problem [13, 19]. For ease of reference we refer to our approach as Bayesian/MaxEnt (BME) reweighting. Note that equivalent or similar reweighting schemes have been used to construct conformational ensembles in biomolecular contexts [20–26].

We begin by briefly describing the underlying theoretical problem, and then exemplify the procedure on a two-dimensional toy model. We then proceed with providing a step-by-step guide for two examples showing how to combine (i) NMR data with MD simulations of a single-stranded RNA tetranucleotide, and (ii) SAXS data with both atomistic and coarse grained simulations of a dynamic, 2-domain protein. In these examples we also show how to use the optimized weights to calculate other structural properties, thereby providing a more accurate description of the system of interest. Throughout the examples we use our software called BME, which is freely available at https://github.com/sbottaro/BME under the GNU GPLv3 license, and where the reader may also find detailed step-by-step guides and examples.

## 2 Methods

### 2.1 Theoretical background

We consider the case in which one samples molecular conformations, x, from a prior distribution *P*^0^(x). Sampling can be performed using e.g. MD simulation with an atomistic force field or Monte Carlo simulations with a coarse-grained model. In practice, our model *P*^0^ is only an approximation of the ‘true’ (but generally unknown) probability distribution *P*^TRUE^. Depending on the system, *P*^TRUE^ may be characterized by a single dominant state (as in a structured protein) or by several distinct states with different populations, as in single stranded RNA or intrinsically disordered proteins. Because of model inaccuracies, *P*^0^ and *P*^TRUE^ may differ: In these cases averages calculated from simulations <*F^calc^*> do not agree with corresponding experimental measurements *F^exp^*.

In the BME approach, one seeks a new probability distribution *P*\* with the following properties:

- It maximizes the relative Shannon entropy:

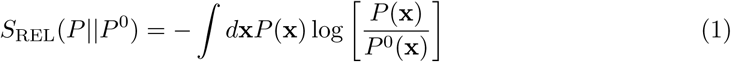
- It matches a set experimental *m* constraints 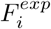 within a tolerance, *ε_i_*, determined via some error model (see further below):

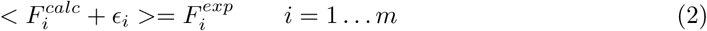
- It is normalized:

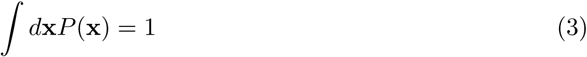

Note that the relative entropy *S*_REL_(*P*||*P*^0^) is the negative Kullback-Leibler divergence [7, 27, 28]: The probability distribution that maximizes the relative entropy can then be considered as the smallest modification to *P*^0^, where the notion of distance in probability distribution space is given by the Kullback-Leibler divergence. A direct link to Bayesian statistics is provided by the observation that the Maximum Entropy distribution is the most probable probability distribution compatible with the data [29].

Here, we consider the discrete case where a finite number of configurations x_1_ … x_*n*_ have been sampled from the prior distribution *P*^0^. The integrals in Eq. 1-3 can then be written as summations over the *n* configurations with corresponding weights 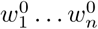, so that 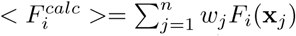.

For methods such as standard MD or MC simulations that generate samples directly from the Boltzmann distribution defined by the force field, the initial weights are uniform 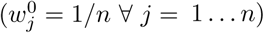. When using biasing techniques such as umbrella sampling [30] or metadynamics [31] they are non-uniform, and have to be estimated using standard techniques prior to using BME.

It can be shown [7, 12, 13, 28] that the weights 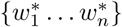 that satisfy Eqs. 1–3 are given by

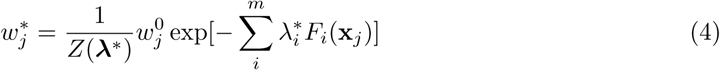

Where the normalization *Z* is defined as

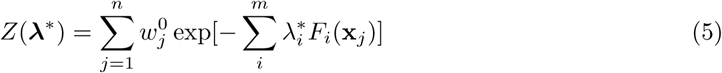

and 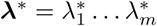 is a set of Lagrange multipliers (one per experimental constraint). When assuming that uncertainties are modeled by independent Gaussian distributions, i.e.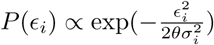the Lagrange multipliers are determined by minimizing the following function [13, 28]:

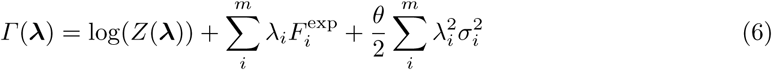

Here, *σ_i_* is the uncertainty on the constraint 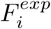 and includes experimental errors and inaccuracies introduced by the calculation of the experimental quantity from a structure (i.e. the forward model).

Because this combined uncertainty is not always known accurately, a global scaling parameter, *θ*, is introduced [17]. When *θ* is large, all *σ* are multiplied by a large factor, and in the limit *θ* →∞ this corresponds to no confidence in the experimental data and reverts to the prior distribution. Conversely, a perfect match with experimental data is achieved when *θ* = 0. Note that when *σ* =0 Eq. 6 reduces to the maximum-entropy solution with no error treatment.

It has been shown that the optimal weights 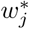 obtained in this way correspond to the weights that minimize the function [17, 10]:

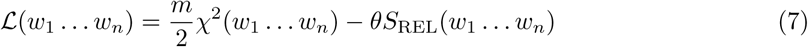

In this equation, χ^2^ quantifies the agreement with the experiments

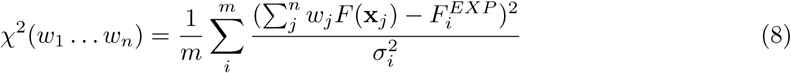

and the relative entropy term 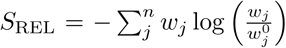 measures the deviation from the initial weights *w*^0^.

Few items are worth highlighting. First, the function 𝓛 in Eq. 7 can be interpreted as a ‘pseudo free-energy’, where χ^2^ plays the role of enthalpy, SREL is the entropy and the parameter *θ* is the temperature. At high temperature (large *θ*) the entropy dominates, while in the limit *θ* → 0 all that matters is to minimize the deviation between experiments and simulation.

While Eq. 7 can more easily be interpreted, it may in practice be difficult to minimize if the number of weights, determined by the number of conformations *n*, is large. One approach previously employed has thus been to cluster the conformations prior to reweighting, thus reducing the number of weights that need to be determined, but at the same time also loosing details present in the original ensemble. Since the Bayesian and Maximum Entropy with error formulations are mathematically equivalent, it is thus in many cases more convenient to minimize Γ (λ) in Eq. 6 rather than Eq. 7, because the number of experimental measurements, *m*, is typically much smaller than the number of frames, *n*.

## 3 Results

### 3.1 Toy model

We illustrate the application and outcome of the above-described reweighting procedure on a twodimensional toy model (Fig. 2A). We construct a model with three states (S1, S2, S3) defined by *P*^TRUE^(*x, y*), that here represents the ‘true’ probability distribution (shown in grey/black). We then assume that it is possible to measure, with some error, the average of the *x* and *y* coordinates, shown as a dark purple square with error bars in Fig. 2A. We then construct a prior distribution *P*^0^(*x, y*) that has the same three states as *P*^TRUE^, but with different populations (red lines). *P*^0^ corresponds, for example, to a situation in which the molecular force field is inaccurate or when sampling is not converged. We sample *n* = 10^5^ (*x, y*) coordinates from *P*^0^, and calculate the average position 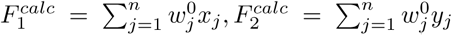. By construction, the calculated averages (red star) are not identical to the ‘true’ averages (black sphere), and the differences to the experimental estimate of these average are greater than the ‘experimental’ error.

**Fig. 2.**
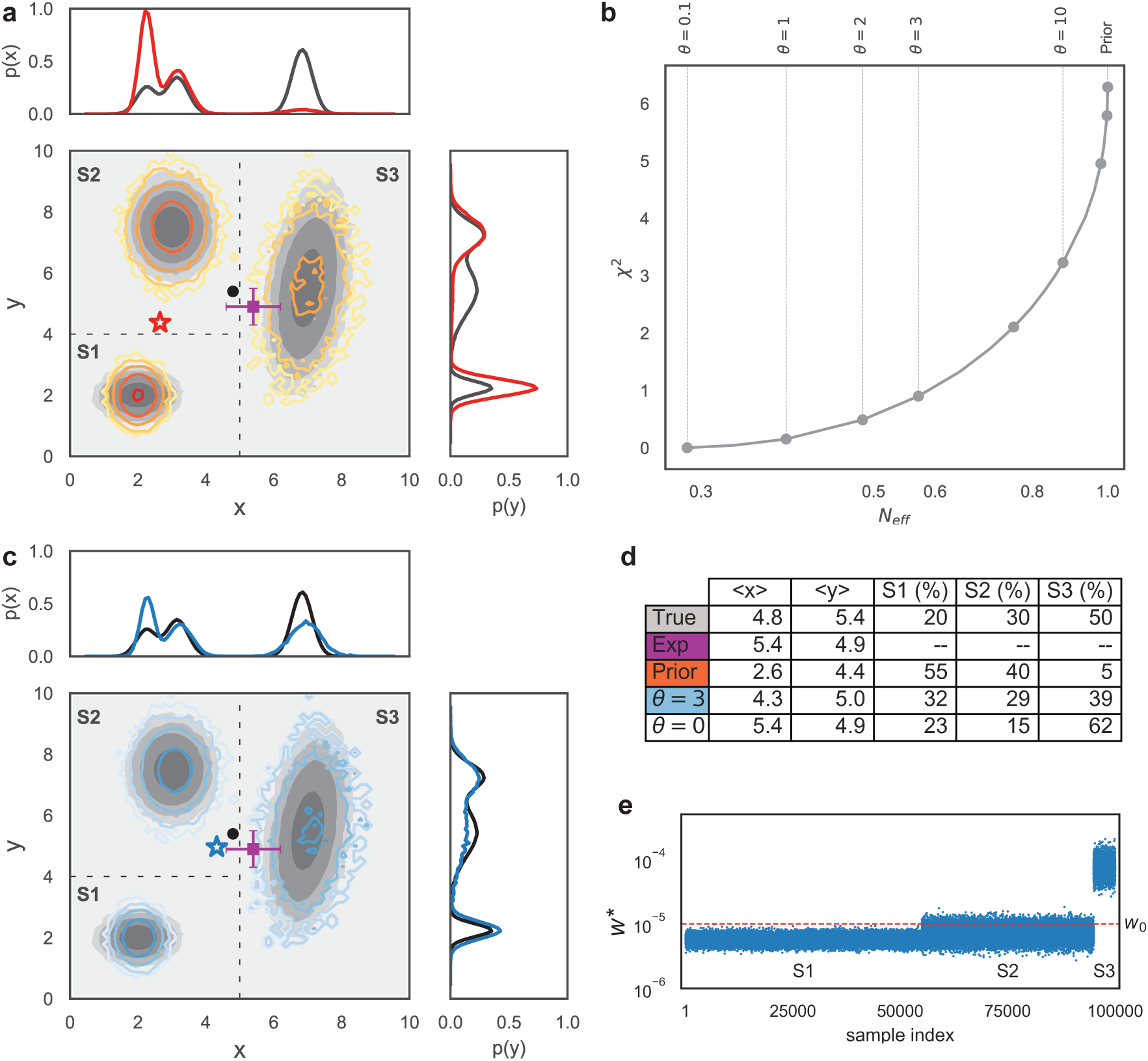
BME reweighting illustrated using a 2D model. (a) *P*^TRUE^(*x, y*) (grey scale) and prior distribution *P*^0^(*x, y*) (orange/red). Both distributions are characterized by three states (S1, S2, S3), but with different populations. The boundaries between the states are shown as dashed lines and are used to calculate the probabilities for the three macrostates, but are not needed in the reweighting analysis. The marginal distributions along x and y are shown as black or red lines. The average *x*, *y* position calculated from P ^0^ (red star) is not compatible with the one calculated from *P*^TRUE^ (black dot). In this example, the dark purple square shows an hypothetical experimental measurement of 〈*x*〉, 〈*y*〉 with associated uncertainty. (b) The effective fraction of frames left after reweighting is shown versus χ^2^ for different values of the parameter *θ*. In this case χ ≈ 1 is obtained using *θ* =3. (c) The histogram calculated using the optimized weights *w*\* with *θ* = 3 is shown in blue and overlaid on *P*^TRUE^ (grey scale). The new average (blue star) is, per construction, in better agreement with the experimentally-measured average position, and also closer to the ‘true’ average. (d) Table reporting the average *x, y* position and the population of the three states S1,S2,S3 for the ‘true’ model, calculated from the prior *P*^0^, and after reweighting using *θ* = 3 and *θ* = 0. (e) Values of the optimized weights *w*\*. The state corresponding to the weights is indicated in the labels, and for visualization purposes the samples were sorted so that samples from the same state are shown together.

Given the initial weights *w*_0_ =1/*n* of each sample, and the ‘experimentally’ measured values and uncertainties, we minimize Γ (λ1,λ2) (Eq. 6) and find the optimal weights *w*\* defined via Eq. 4. This procedure is repeated for different values of *θ*. At high values of *θ*, the entropy term dominates and the weights are close to their initial (uniform) values. As *θ* is decreased, the weights become less uniform as they are reweighted to find a combination to match the experiment better (decrease χ^2^). This decrease in ‘flatness’ of the weights corresponds to a drop in the number of frames that effectively contribute to the calculated averages, and can be quantified by the ‘effective fraction of frames’ *N_eff_* = exp (*S*_rel_). We thus find it useful to plot Neff versus χ^2^ to illustrate the balance between the requirement of fitting the data well (low χ^2^) and minimally perturbing the prior distribution (large *N_eff_*) (Fig. 2B).

Inspection of this plot shows as expected that when *θ* → 0 we achieve a very good agreement between simulation and ‘experiments’ (χ^2^ → 0). At the same time, we introduce a large perturbation to the prior probability distribution *P*^0^, so that the relative entropy is a large, negative number and the fraction of effective frames becomes small. In the limit of large *θ*, instead, χ^2^ approaches the initial value obtained when sampling from *P*^0^, the new weights *w*\* are close to wand thus the number of effective frames *N_eff_* approaches 1. A practical solution to the trade-off between the two limits can be found by scanning different values of the parameter, starting from a large number, until a further decrease in *θ* does not result a significant decrease in the associated χ^2^. Such a procedure, often termed finding the ‘elbow’ of the curve (and similar to L-curve selection in other regularization techniques), provides a range of viable values for *θ*: in the toy model, for example, one could pick *θ* = 3, that leads to a χ^2^ ≈ 1 for both x and y coordinates. After fixing *θ* = 3 we can observe the modification to the probability distribution introduced by reweighting (Fig. 2C). First, the calculated average (blue star) is close to the experimental average (within a level set by the uncertainty). Additionally, the reweighted (blue) and ‘true’ distribution (grey) are similar one to another.

In this toy model we have three distinct states, whose population can be calculated by summing the weights of the samples belonging to each region. In Table 2E we report these population for the ‘true’ model, for the prior distribution *P*^0^ and after reweighting with *θ* = 3 and *θ* = 0. We note that this kind of clustering into states is not a part of the actual reweighting procedure, but may be useful in the subsequent analyses. We can see that the population of state S1 decreases from 55% in the unreweighted ensemble to ≈ 25% in the reweighted one. The population of S3, instead, is increased from 5% to 40%, substantially closer to the ‘true’ population of 50%. Note that it is in principle possible to obtain a better agreement with experiments by setting *θ* = 0 (Table 2D). In a realistic scenario this would not be advisable, since experimental quantities are not known with infinite precision, and setting *θ* = 0 effectively corresponds to ignoring the uncertainties in the experiments and forward models.

Finally, it is instructive to plot the individual weights *w*\* for each sample (Fig. 2E). In agreement with the population shift described above, we can see that all samples belonging to state S1 are down-weighted with respect to the initial weights *w*^0^, while the opposite effect happens to samples belonging to S3.

As is clear from the example above, the BME reweighting procedure enables the reconstruction of an ensemble that is closer to the ‘true’ ensemble by combining the imperfect prior model with the experimental data. Before proceeding to discuss application in molecular simulations and structural biology, we note, however, that there might occur situations in which the reweighting approach described here would provide incomplete or wrong result. More precisely, we identify four possible sources of problems:

- *Non-informative experimental data*. There might be situations in which the available experimental data cannot substantially correct the inaccuracies of the model. In the toy model, this would correspond for example to knowing the average *y* position but not *x*. In such a situation, the ‘true’ population of state S3 could not be determined very accurately because the average *x* position carries information on the relative populations of S1+S2 with respect to S3. In practice for high-dimensional systems such as biomolecular ensembles, the situation is more complex. Indeed, most experimental measurements are sensitive to some, but not other aspects of the distribution of conformations, and generally there are many more degrees of freedom than experimental observations. Indeed, it is this underdeterminism that necessitates the use of the prior model.
- *Insufficient sampling*. If sampling is not exhaustive, relevant states are not explored, and thus it is not possible to estimate with any certainty their weights after reweighting. This observation is equivalent to the well-known problem of large uncertainties in estimating free energies between states with little overlap. In such cases a small *θ* (corresponding to a small *N_eff_*) could be required to achieve a reasonable agreement between simulations and experiments. As a consequence, most of the optimized weights are vanishingly small, and a small set of weights dominates the ensemble. In this situation, longer simulations or the use of enhanced sampling techniques are necessary, and could e.g. be guided by the structures whose weights are increased at intermediate values of *θ*. In practice, the prior that enters the reweighting approach is not the full distribution, *P*^0^, but rather our estimate of this from the finite samples representing the starting ensemble.
- *Inaccurate force field*. The reweighting approach relies on the accuracy of the prior distribution, in particular when the data is sparse and noisy. One can imagine for example the case of a uniform prior distribution over *x, y* (within some range) in the toy model. A small modification to this distribution would lead to a very good agreement with the experimental data, but would not be close to the ‘true’ probability distribution. Indeed, the BME formalism is not guaranteed to give the best model possible. Instead, it provides the least biased model that takes into account all the knowledge we have of the system, encoded both in the (potentially inaccurate) force field and the (noisy and sparse) experimental data, and no more than this information. If more information were available or assumed (such as assuming that the ensemble is narrow), this should preferably be encoded and input into the model.
- *Inconsistent or wrong experimental data*. When data are inconsistent, the reweighting approach might fail in obtaining improved agreement with all experimental data. For example, in our previous work we have identified spectral overlaps by noticing that a subset of NOE distances could not be reweighted [19]. In certain cases, the presence of inconsistent data points could be detected via cross-validation. In general, however, there is no guarantee that erroneous data cannot be fitted with an erroneous ensemble, and indeed ensemble fitting can be prone to such ‘overfitting’ to erroneous data. When very small values of *θ* are needed to fit the data, this can be a sign of either a poor prior, underestimation of the uncertainty in the data, or actual errors in the experiments.

In realistic cases, these problems can occur simultaneously, and can be sometimes difficult to disentangle. We also stress that BME is inherently an ensemble refinement procedure [17], and the successful use of this approach depends on the amount/quality of experimental data, on sampling and on force field accuracy. Further discussions of problematic situations in ME approaches are also found in Refs [10, 28, 6].

### 3.2 Combining NMR data with MD simulation of RNA

Following the above introduction of the BME method with the two-dimensional toy model, we now proceed to describe how to obtain the conformational ensemble of an RNA tetranucleotide using atomistic MD simulations in combination with experimental data from nuclear magnetic resonance (NMR) spectroscopy. In particular, we describe how to carry out the analysis and suggest some best practices using the Python code for BME that is freely available under under a GNU GPLv3 license license at https://github.com/sbottaro/BME. Data and a detailed list of commands to perform the analysis described below are available on the github repository.

In essence, the procedure can be summarized in four steps: 1. Data collection and preparation, 2. Minimizing Γ and parameter selection 3. Cross validation and 4. Interpretation of the weights and of the reweighted ensembles.

**Step 1: Data collection and preparation** The first step is to collect and format experimental and simulation data necessary for BME reweighting. The experimental datafile(s) contains *m* experimental averages and uncertainties, and has the following format:

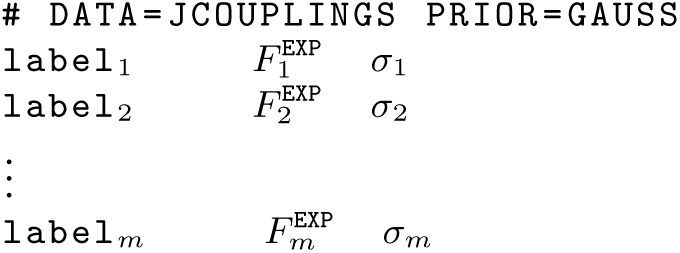

The first line is a header that specifies the type of input data and the type of prior on the error. Currently, the BME software specifically supports the following data types: nuclear Overhauser effect (NOE), scalar couplings (JCOUPLINGS), chemical shifts (CS), small-angle x-ray scattering (SAXS), and generic distance restraints (DIST). Only a Gaussian (GAUSS) error prior is implemented in the current version of the BME software. Since all-linearly average data are treated in the same way, other sources of information can use these functions. The first column is a user-defined label that indicates e.g. the torsion angle in a ^3^J scalar coupling or the name of the protons involved in the NOE. The second column is the experimental value and the third the associated uncertainty. For NOEs, it is possible to specify upper/lower boundaries instead of average values. In such cases, the restraint is applied only if 〈*F^calc^*〉 is larger/smaller than *F^EXP^*. This information can be specified by flagging the experimental datafile in the following fashion:

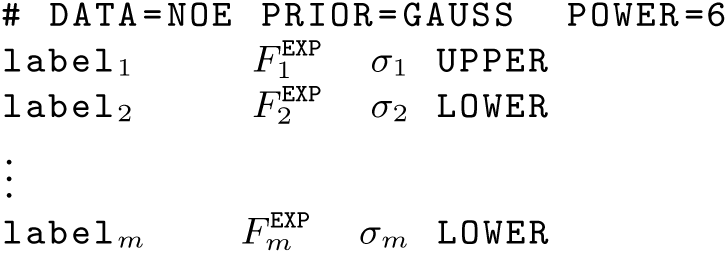

For most types of experiments, the underlying model is the same in the sense that the ensemble averaged values is simply a linearly-weighted average over the values calculated for each frame, e.g. for scalar couplings 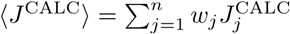. When using NOE data, however, the experimental data has to be specified as a distance *r*^EXP^, and the imposed restraint is proportional to the volume of the corresponding peak in a NOESY spectrum, i.e. 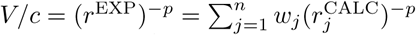. The power p, is by default set to 6, but it can be set to a different value (e.g. 3 [32]) using the keyword POWER in the header of the experimental datafile.

The other information required for reweighting are the values of the experimental observables calculated for each frame in the simulation. The data is stored in a file with the following format

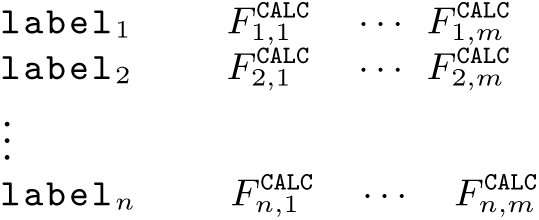

Here, 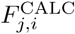 corresponds to the *i*^th^ experimental average (as ordered in the experimental datafile) calculated on the *j*^th^ frame. The label is user-defined and can for example be the frame number. The number of frames *n* can be on the order of tens or hundreds of thousands: there is in principle no restrictions on *n* since the complexity of the problem is mostly determined by the number of experimental restraints *m*. Note also that the back-calculation is performed only once, and any type of forward model can be used as long as the calculated values can be written as a weighted average over the input configurations.

As an example, we here consider experimentally measured NMR ^3^J scalar couplings in an RNA tetranucleotide [33] and extensive MD simulation of the same system taken from our previous study [19]. In Fig. 3a we show in grey the *m* =25 experimental measurements, sorted by magnitude for visualization purposes. The averages calculated directly from the simulations using *n* = 20000 frames are shown in red. Simulations and experiments do not agree perfectly: in this case at least five MD averages do not fall within experimental uncertainty.

**Step 2: Γ minimization and *θ* selection** Given the data described above, and at a fixed value of *θ*, we minimize the function Γ defined in Eq. 6 with respect to the *m* Lagrange multipliers. In our implementation, we use the limited memory Broyden-Fletcher-Goldfarb-Shanno (L-BFGS) optimization algorithm implemented in the NumPy library. Since the analytic gradient can be easily calculated, the optimization is computationally inexpensive and typically takes seconds on a standard desktop computer with *m* ≈ 10^2^ and *n* ≈ 10^4^ − 10^5^. The optimization returns a set of optimal λ that are used to calculated a weight for each frame via Eq. 4. By definition, the optimal weights improve the agreement with input experimental data. As described in the previous section, we then scan different values of the parameter *θ*, and consider how the number of effective frames *N_eff_* and χ^2^ vary as a function of this parameter.

In our RNA tetranucleotide example (Fig. 3b), a small value of *θ* corresponds to a better fit with scalar couplings (low χ^2^), but obtained at the cost of a large drop in relative entropy and hence only a small effective fraction of frames used. Using a large *θ*, instead, we approach the χ^2^ of the prior distribution. In this case one can identify useful values of *θ* in the range 2 − 10, corresponding to the ‘*elbow*’ region in the *N_eff_* versus χ^2^ plot.

**Step 3: Cross-validation** When possible, it is recommended (at least initially) to split the experimental data into some used in optimization and some used for cross-validation. In our example, we monitor the behavior of the NOE data, that were not used as input for reweighting. The agreement with NOE distances has a clear minimum around *θ* = 10. Note that when *θ* =0.1 the χ^2^ relative to the NOEs is high, meaning that enforcing a tight agreement with scalar couplings is detrimental. Here, we choose *θ* = 2 as it provides improved agreement with scalar couplings without a dramatic drop in *n_eff_*. Having fixed *θ* = 2, it is possible to calculate the individual NOE averages before (red) and after (blue) optimization (Fig. 3c).

**Fig. 3.**
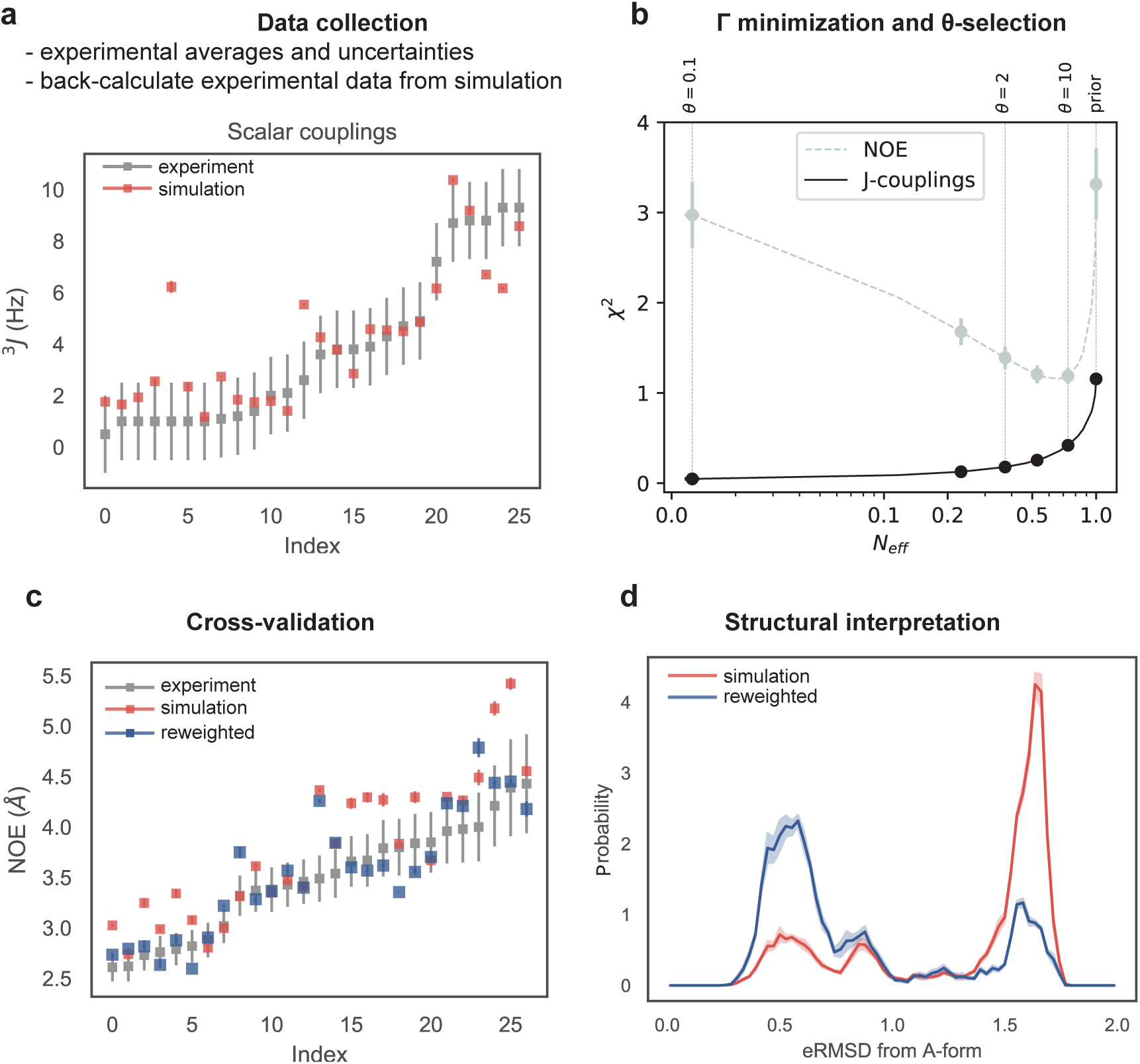
Experimentally-restrained simulation of an RNA tetranucleotide. a: Experimental ^3^J couplings (grey) compared to calculated averages using the original MD simulation (prior distribution, shown in red). Error bars indicate experimental uncertainties (grey bars) or the standard error of the mean estimated using 5 blocks (red bars; typically, smaller than the point). b: *N_eff_* versus χ^2^ plot using scalar couplings restraints for different values of *θ*, and cross-validation using NOEs. c: Experimental NOEs (grey) compared to calculated averages using the original MD simulation (red) and after reweighting using scalar couplings data (blue). d: Histogram of the eRMSD from an A-form RNA structure calculated from the original MD simulation (red) and after reweighting (blue).

**Step 4: Structural interpretation** In order to understand how the new weights affect the original MD conformational ensemble, it is useful to calculate the distribution of various structural parameters. As an example we show the histogram of the eRMSD [34, 35] from a reference A-form helix of the original MD simulation (prior) and of the restrained one (Fig. 3d). The effect of the experimental restraint is to favor the presence of structures closer to A-form, as also described in our previous work [19].

### 3.3 Combining SAXS data with simulation of proteins

We now proceed to apply the BME method and software on a larger, more complex system with less informative experimental data. In particular, we show the results for a two-domain protein that shows substantial structural heterogeneity of the relative orientation of the two domains, and describe how we can refine our molecular simulations by reweighting against small-angle X-ray scattering (SAXS) data. The reweighted ensemble in turn allows us to improve our description of the dynamical domain-domain motions. As object for our study we chose the protein sf3636 from the bacterium *Shigella flexneri 2a*. Previous studies have shown that the protein consists of two structural domains (NTD and CTD), with substantial interdomain motions as probed via different experiments including SAXS [36], thus providing a good example for applying BME to proteins.

Because large-scale motions in proteins might be difficult to sample with conventional simulations, they are attractive targets for coarse-grained (CG) simulations. Specifically, we here applied a recently updated parameterization of the Martini CG model [37]. In line with standard recommendations for studying protein dynamics with Martini, we applied harmonic restraints to keep each of the folded domains relatively rigid [38], whereas no restraints were applied between pairs of atoms spanning between the NTD and CTD. We performed a 4 μs long simulation using MARTINI 3.0beta with the force constant for the harmonic restraint set to the default 500 kJ/(mol· nm^2^) using the Gromacs software [39] and standard settings. We analysed and reweighted an ensemble consisting of 8000 structures from the simulation by taking each 500 ps frame (Fig. 4a).

**Fig. 4.**
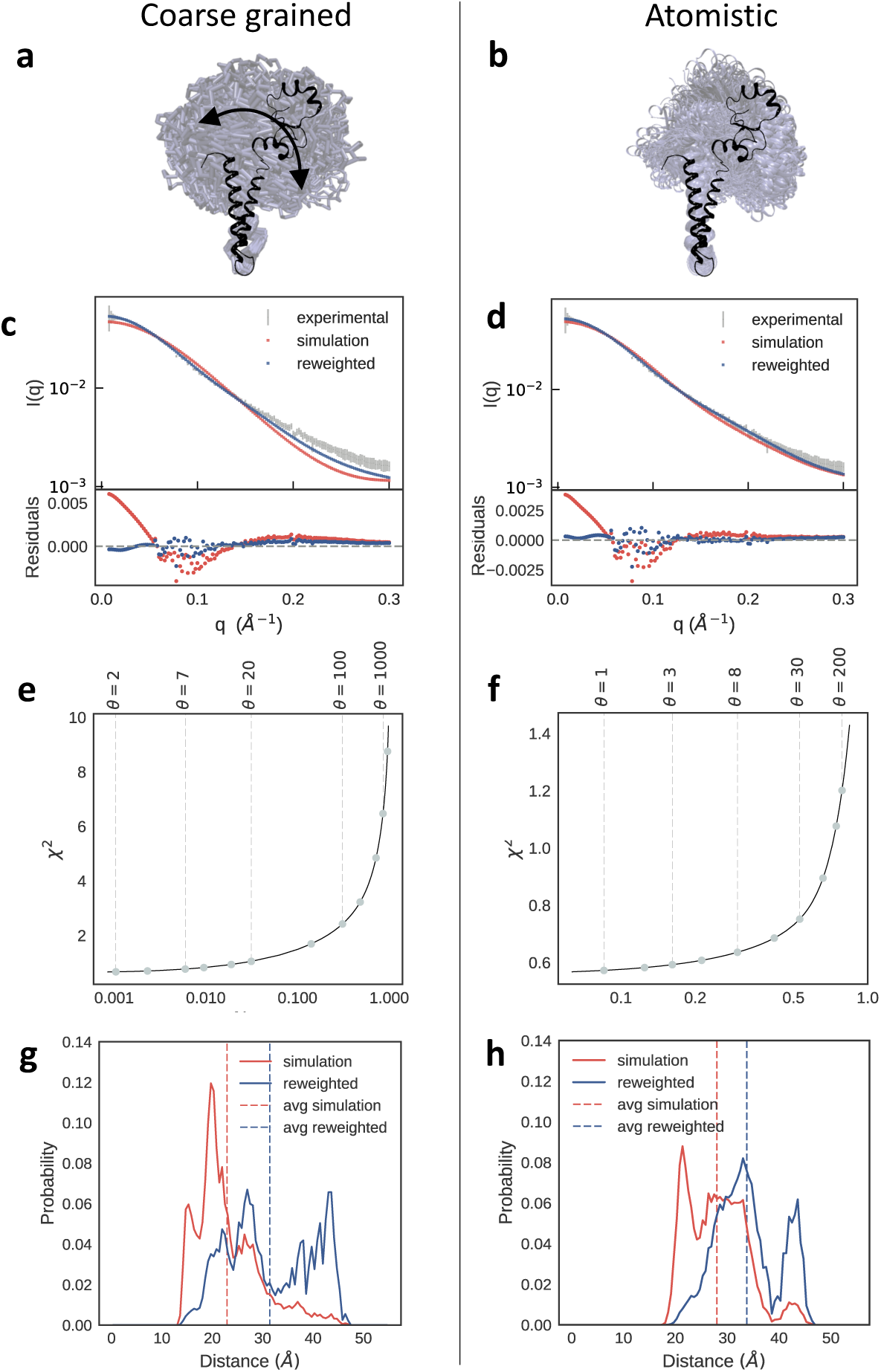
SAXS refinement of coarse-grained and atomistic simulations of a flexible two-domain protein. Left (a,c,e,g), refinement of a coarse-grained MARTINI 3.0b simulation with an elastic network force constant of 500 kJ/(mol·nm ^2^). Right (b,d,f,h), refinement of an atomistic simulation using the a99SB*disp* force field. (a,b) The black structure is the starting configuration, and the blurred blue illustrates the sampled configurational space for each simulation by showing every fifth frame in the simulation aligned to the (bottom) NTD domain. (c,d) Calculation of SAXS data from the original MD simulations and the refined ensembles are compared to the experimental data. (e,f) Evaluating the effect of the global scaling parameter *θ* to balance the prior (force field) and the experimental data. For the atomistic simulation we chose *θ* = 30, and for the coarse grained simulation we chose *θ* = 100. (g,h) Analysis of the effect of reweighting against experimental data on the interdomain distance.

For comparison, we also performed an all-atom, explicit solvent simulation using the a99SB**disp** force field [40]. This force field has recently been parameterized to provide an accurate balance between protein-protein and protein-water interactions, and thus should be particularly useful for looking at transient interactions between the two domains. Specifically, we performed a 2 μs long simulation using a time step of 2 fs, a temperature of 298 K and 1 bar pressure with the velocity rescaling thermostat [41] and Parrinello-Rahman barostat [42]. We analysed and reweighted an ensemble consisting of 20,000 structures from this simulations by taking one frame every 100 ps in the simulation (Fig. 4b).

We used Pepsi-SAXS [43] to calculate the SAXS data for each of the extracted structures from the two simulations (Figs. 4c and 4d). In the case of the Martini simulation we first used standard approaches to reconstruct all-atom models from the CG beads [44]. As Pepsi-SAXS has several free parameters whose values may vary between proteins, we estimated these values from the ensembles. Because optimizing the values from each conformation might lead to substantial overfitting, we instead used an approach where we (for each ensemble) used the average value of such-optimized Pepsi-SAXS parameters over the entire ensemble and reran Pepsi-SAXS with these parameters fixed. We then compared the ensemble-averaged SAXS data from the atomistic and the coarse grained ensembles with previously determined experimental values [36]. The results show that while both simulations are in reasonably good agreement with experiments, the all-atom simulations appear provide a better description of the structure and dynamics in sf3636 (Figs. 4c and 4d).

Despite the good overall agreement, both simulations show systematic discrepancies with experiments, in particular at low-to-middle values of q. We thus use the BME procedure to refine the simulations of sf3636 against the SAXS data. Here, the experimental input file and the simulation input file with SAXS calculations for each frame in the ensemble have the following formats:

Experimental file format: Simulation SAXS file format:

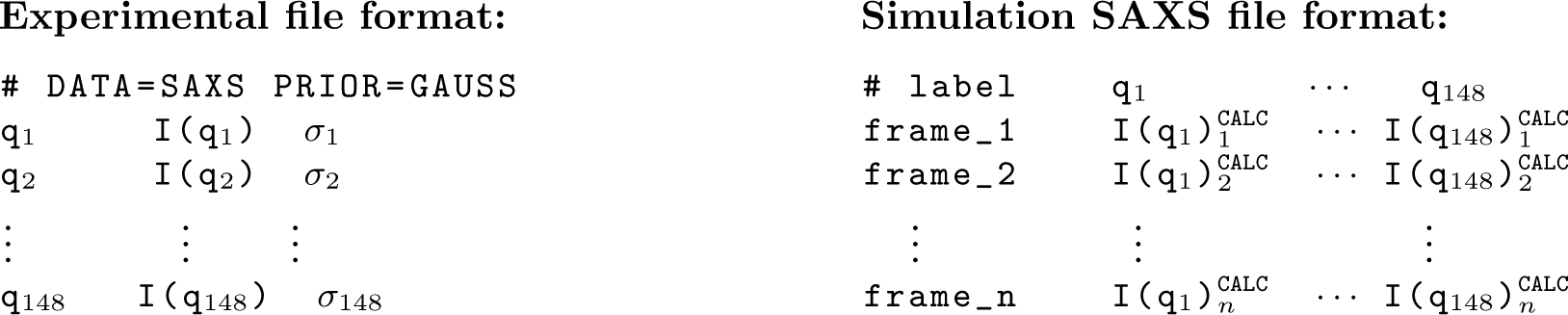

We determine the χ^2^ between the calculated and experimental scattering intensities over all scatter vector points, *q_i_*, in the range 0.006-0.3 Å^−1^. Because of difficulties in estimating errors in scattering intensities and due to correlations between neighbouring points the absolute value of χ^2^ may be difficult to interpret for a SAXS experiment [45]. Thus, for each ensemble we analysed the*N_eff_*-vs-χ^2^ plot to find a value of *θ* that reflects the compromise between the prior (simulation) and the data (Figs. 4e and 4f). From these we choose *θ* = 30 for the all-atom simulation and *θ* = 100 for the coarse grained simulation, and the resulting calculated SAXS curves are, as expected, in much better agreement with experiment (Figs. 4c and 4d).

With the optimized weights it becomes possible to analyse other properties of the conformational ensembles. As an illustration, and following the previous study of this protein [36], we analysed the distribution of the interdomain distance, quantified as the distance between the centres of mass of the NTD and CTD. The resulting histograms show that the reweighting in general has the effect of increasing the interdomain distances, suggesting that despite recent force field improvements for both coarse-grained and all-atom MD simulations they might still overestimate protein-protein interactions. Thus, the BME approach has the potential for making the resulting ensembles more robust than those from the unbiased simulations, thereby removing some of the uncertainty coming from the imperfect force fields.

## 4 Notes

- Our software, BME, is freely available on the Internet at https://github.com/sbottaro/BME together with examples on how it can be used.
- When using biasing enhanced sampling techniques such as umbrella sampling or Metadynamics, the weights of the prior distribution are not uniform, i.e. 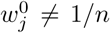. The BME reweighting approach can be applied in this case by using the function `set weights()` before calling the function `optimize()`.
- The BME software can easily be extended to include additional types of experimental data that can be calculated as the average over the weighted contribution from each frame. Experimental data that depend also on global parameters that need to be optimized are currently not supported. Data that depend on temporal correlations (e.g. kinetic data) are also not supported.
- It is advisable to assess the robustness of the procedure by performing several cross-validation tests and by using blocking procedures for error estimation.

## 5 Acknowledgements

We thank Dr. Alexander Lemak and Prof. Cheryl H. Arrowsmith for sharing the SAXS data on sf3636. We also thank Yong Wang and Mustapha Carab for input to and testing of BME. The research and development described here were supported by a grant from The Velux Foundations, a Hallas-Møller Stipend from the Novo Nordisk Foundation, and the Lundbeck Foundation BRAINSTRUC initiative.

